# Role of Paravertebral muscle myostatin upregulation in the development of idiopathic scoliosis

**DOI:** 10.1101/2022.05.23.493176

**Authors:** Jiong Li, Gang Xiang, Sihan He, Guanteng Yang, Chaofeng Guo, Mingxing Tang, Hongqi Zhang

## Abstract

Paravertebral muscle (PVM) abnormalities play important roles in the pathogenesis of idiopathic scoliosis (IS), and elevated oxidative stress could result in PVM injury in IS patients, but the underlying mechanism of oxidative stress generation is still unclear. Increased apoptosis, impaired myogenesis and elevated oxidative stress were found in primary skeletal muscle mesenchymal progenitor cells (hSM-MPCs), which are essential for the myogenesis process of vertebrate skeletal muscles, of IS patients. Through RNA-sequencing and further analysis, we identified significantly upregulated myostatin (MSTN) in IS hSM-MPCs. Overexpression of MSTN in hSM-MPCs from control patients increased the expression of NADPH oxidase 4, promoted reactive oxygen species production and apoptosis, and suppressed myogenesis. However, MSTN knockdown decreased the expression of NADPH oxidase 4, inhibited reactive oxygen species production and apoptosis, and enhanced myogenesis in IS hSM-MPCs. In addition, overexpression of MSTN in the PVMs of mice induced elevated oxidative stress and scoliosis without abnormal vertebral structure. Altogether, our study suggested that abnormal PVM changes and accumulated oxidative stress in IS patients may result from upregulation of MSTN, which could contribute to the development of IS.

## Introduction

Idiopathic scoliosis (IS) is a three-dimensional spinal deformity, the peak incidence of which is around puberty (namely, adolescent idiopathic scoliosis, AIS). Since the important questions concerning its etiology and pathogenesis remain poorly understood (Fadzan & Bettany-Saltikov, 2017), treatment of IS involves only long-term bracing and costly surgical prevention (Zhuang *et al*, 2019). The former has limited effectiveness, while the latter has high risk. Therefore, the need to reveal the pathogenesis of IS development and discover effective curative strategies is urgent.

Spinal stability is influenced not only by the spinal column but also by the paravertebral muscle (PVM); thus, the idea that abnormalities in the PVM might be the cause of idiopathic scoliosis has been entertained for decades (Lowe *et al*, 2000). Extensive studies have shown severe pathological changes in the PVMs of patients with IS. Disrupted myofibrillar elements, muscular atrophy due to necrosis, infiltration of immune cells and adipocytes, and the presence of hyaline fibers were also found in the PVM of AIS patients (Jiang *et al*, 2017; Khosla *et al*, 1980; Wajchenberg *et al*, 2015). Our previous study similarly noticed increased myofiber necrosis and apoptosis and an altered percentage distribution of myofiber types in IS patients compared with those in matched control subjects. Furthermore, an important role of elevated oxidative stress has been newly shown in these abnormalities of the PVM from IS patients (Li *et al*, 2019a). However, the reason why oxidative stress is increased in the PVM of IS patients is still unclear.

In addition, with the development and progression of scoliosis, concave and convex sides emerge. On the one hand, observed differences in fiber types and degrees of fibrosis and fatty involution between the PVM of the concave side and the convex side suggest that the imbalance of the concave and convex sides may lead to the development of IS (Lleras-Forero *et al*, 2020; Luo *et al*, 2022; Shao *et al*, 2020). However, on the other hand, other scholars reported no significant differences between the concave and convex sides, and the onset of IS was considered to be related more to changes in muscle structure and function rather than differences resulting from the occurrence of IS (Li *et al.,* 2019a; Ng *et al*, 2022; Yeung *et al*, 2019). Nevertheless, the stress that muscles endure from the concave and convex sides differs once the curve is formed, and the differences between the two sides that are generated after the formation of the curve or that lead to the development of deformity are uncertain.

Because human skeletal muscle mesenchymal progenitor cells (hSM-MPCs) play important roles in the myogenesis process of vertebrate skeletal muscles and elevated oxidative stress can induce myofiber injury in hSM-MPCs, in this study, primary hSM-MPCs from the PVM of IS patients and controls were isolated for further examination of the mechanism of elevated oxidative stress and abnormal changes in the PVM of IS patients (Buckingham, 2001; Li *et al.,* 2019a). The relationship between abnormal changes in the PVM, including concave-convex side differences, and the development of IS was investigated in vivo.

## Results

### General physiological characteristics of IS patients and control groups

The PVM tissues of 15 controls and 15 IS patients were used for primary hSM-MPC extraction in our study. The mean age of the IS patients was 17.53 years, while the mean age of the control patients was 18.40 years, with no significant difference. There were 8 females/7 males in the control group and 10 females/5 males in the IS patient group, and sex differences between the two groups were not observed according to a *x*^2^ test (Table 1 and Table EV1). In addition, the average Cobb angle of the major curve in the IS patients was 47.78° (Table 1).

**Table 1.** Demographic of study populations.

| Characteristics | CT *N*=15 | IS *N*=15 | *P* value |
| --- | --- | --- | --- |
| Age (years) | 18.40 ± 3.64 | 17.53 ± 3.42 | 0.507 |
| Gender | F 8 / M 7 | F 10 / M 5 | 0.710 |
| Major curve Cobb Angle (°) | — | 47.78 ± 13.84 | — |

CT, control group; IS, idiopathic scoliosis patients; F, female; M, male. Data was present as Mean ±
Standard Deviation.

### Increased apoptosis, impaired myogenesis and elevated oxidative stress in hSM-MPCs from IS patients

Given that increased apoptosis and abnormal myogenesis in the PVMs of IS patients were observed, we also examined the changes in apoptosis and myogenesis between hSM-MPCs from IS patients (IS-hSM-MPCs) and controls (CT-hSM-MPCs). As shown in Fig 1A and B, the TUNEL assay showed that the proportion of TUNEL-positive cells was significantly increased in IS-hSM-MPCs. The protein levels of BCL2-associated X protein (Bax) and the Bax/B-cell leukemia/lymphoma 2 (Bcl2) ratio in IS-hSM-MPCs were significantly higher than those in CT-hSM-MPCs, while no significant changes in Bcl2 were observed (Fig 1C). Moreover, immunofluorescence staining for the myofiber marker Desmin showed that the myogenesis differentiation ability of IS-hSM-MPCs was weaker than that of CT-hSM-MPCs (Fig 1D). Meanwhile, the protein levels of the myogenesis regulator myogenin (MYOG) in IS-hSM-MPCs were significantly decreased compared with those in CT-hSM-MPCs (Fig 1E). Since our previous study observed elevated oxidative stress in the PVM of IS patients and oxidative stress can induce apoptosis and abnormal myogenesis in hSM-MPCs, we also investigated reactive oxygen species (ROS) production in IS-hSM-MPCs and CT-hSM-MPCs. ROS assay results showed increased ROS production in IS-hSM-MPCs (Fig 1F). However, the protein levels of the oxidative stress-related enzyme endothelial NO synthases (eNOS), which was increased in the PVM of IS patients, were not different between the two cell groups (Fig 1G).

**Figure 1.**
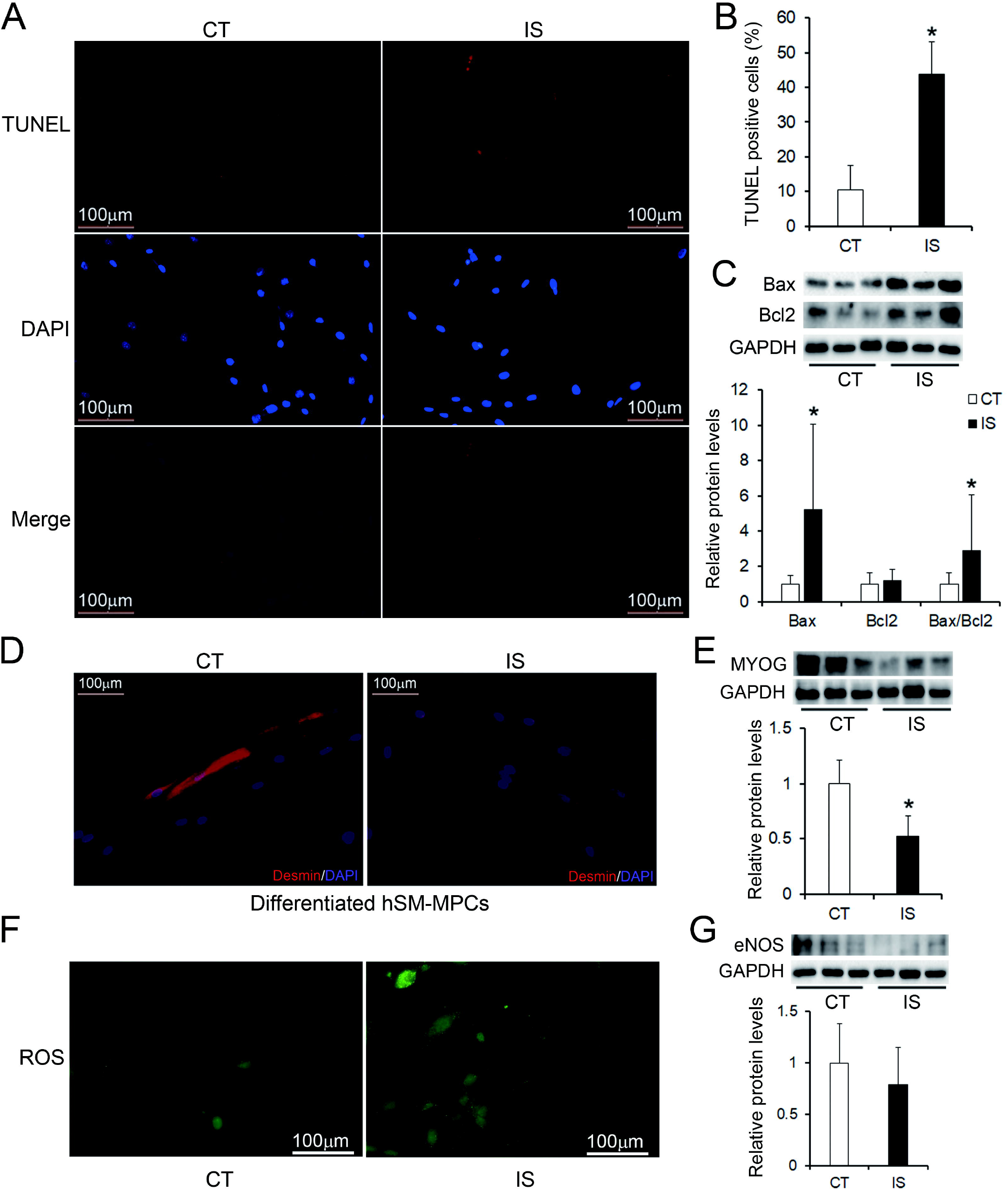
Abnormal changes of IS-hSM-MPCs. A TUNEL stained IS-hSM-MPCs and CT-hSM-MPCs, 200×. B Quantity evaluation of percentage of TUNEL positive cells. C Western blotting analysis of apoptosis related proteins and quantification results in IS-hSM-MPCs and CT-hSM-MPCs. D Desmin/DAPI co-stained IS-hSM-MPCs and CT-hSM-MPCs after differentiation. E Western blotting analysis of myogenesis regulator Myog and quantification results in IS-hSM-MPCs and CT-hSM-MPCs. F ROS production of IS-hSM-MPCs and CT-hSM-MPCs. G Western blotting analysis of ROS related proteins in IS-hSM-MPCs and CT-hSM-MPCs. n=15, *p<0.05. IS-hSM-MPCs, hSM-MPCs from IS patients; CT-hSM-MPCs, hSM-MPCs from controls; ROS, reactive oxygen species.

These results together showed increased apoptosis, impaired myogenesis and elevated oxidative stress in IS-hSM-MPCs, which were similar to the pathological changes in the PVM of IS patients that we reported before. Thus, we speculate that unrelieved oxidative stress further causes apoptosis and a dysregulation of myogenesis in IS-hSM-MPCs.

### Upregulation of MSTN in IS-hSM-MPCs

To determine the underlying mechanisms of the increased ROS production and abnormal changes observed in IS-hSM-MPCs, we conducted RNA-seq analysis of hSM-MPCs from 3 IS patients and 3 controls. A total of 302 genes were identified that differentially expressed (Q value <0.05) between the two groups, among which 153 genes were upregulated and 149 genes were downregulated (Fig 2A and B). Further Kyoto Encyclopedia of Genes and Genomes pathway analysis of the differentially expressed genes showed no significantly associated pathways in our initial analysis (Fig EV1). However, we identified five skeletal muscle development- and oxidative stress-related genes (Fig 2C), and their expression levels were verified by RT-qPCR in 15 IS-hSM-MPCs and 15 CT-hSM-MPCs. As shown in Fig 2D, the mRNA levels of *HIF1A* (hypoxia inducible factor 1 subunit alpha), *PRDX2* (peroxiredoxin 2), *WNT5B* (wingless-type MMTV integration site family, member 5B) and *SOX6* (SRY-box transcription factor 6) were not different between IS-hSM-MPCs and CT-hSM-MPCs, but significantly upregulated mRNA levels of *MSTN* (myostatin) were observed in IS-hSM-MPCs. Increased protein levels of MSTN in IS-hSM-MPCs from IS patients were also validated by western blotting (Fig 2E). We further investigated the mRNA levels of MSTN in the PVMs of IS patients and controls, and significantly increased MSTN was also observed in the PVMs of IS patients (Fig EV2).

**Figure 2.**
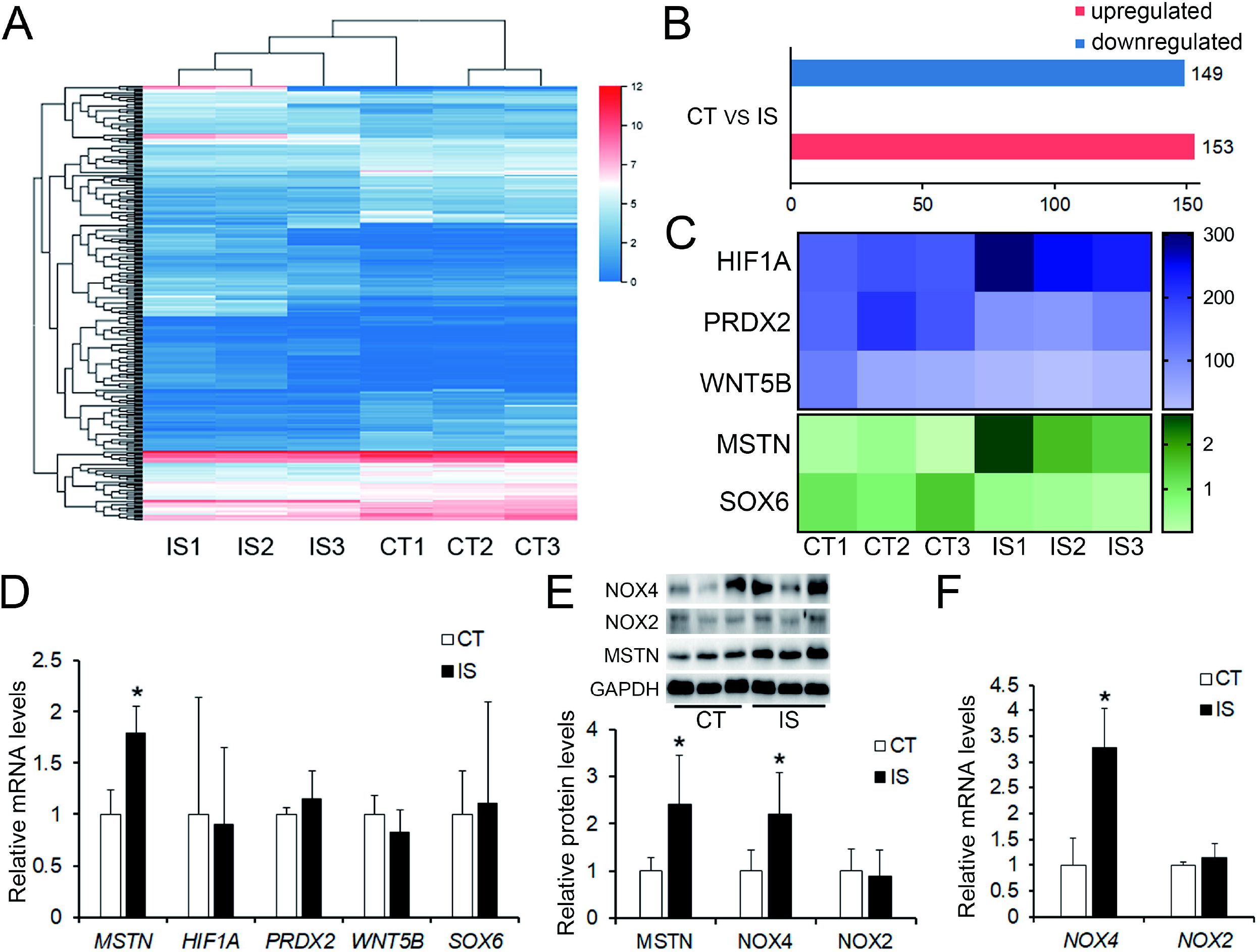
Upregulation of MSTN in IS-hSM-MPCs. A, B Cluster analysis of 302 differentially expressed genes which include 149 down-regulated and 153 up-regulated genes from RNA-seq results between hSM-MPCs of 3 IS patients and 3 control people. C Significantly changed 5 genes associated with skeletal muscle development and oxidative stress according to the RNA-seq results. D The mRNA levels of skeletal muscle development and oxidative stress related 5 genes in IS-hSM-MPCs and CT-hSM-MPCs, n=15, *p<0.05. E Western blotting analysis of MSTN and NADPH oxidases and quantification results in IS-hSM-MPCs and CT-hSM-MPCs, n=15, *p<0.05. F The mRNA levels of NADPH oxidases in IS-hSM-MPCs and CT-hSM-MPCs, n=15, *p<0.05.

MSTN has been shown to inhibit the myogenic differentiation of hSM-MPCs/myoblasts, to function as a negative regulator of skeletal muscle growth and to regulate NADPH oxidase 4 (NOX4) and ROS release (Verzola *et al*, 2020; Zhang *et al*, 2018b). Therefore, we next examined the expression levels of the main NADPH oxidases, NADPH oxidase 2 (NOX2) and NOX4, in hSM-MPCs from IS patients and control group. Compared with CT-hSM-MPCs, significantly increased levels of NOX4 mRNA and protein were found in IS-hSM-MPCs, but there were no significant differences in NOX2 levels between the two cell groups (Fig 2E and F). Our results suggest that MSTN may induce ROS release by upregulating NOX4 in IS patients.

### MSTN overexpression promotes ROS production and apoptosis and suppresses myogenesis in CT-hSM-MPCs

To verify the effect of MSTN upregulation, we overexpressed MSTN in CT-hSM-MPCs (Fig 3A and B). As shown in Fig 3C, MSTN overexpression increased the protein levels of Bax and the Bax/Bcl2 ratio but not Bcl2. The proportion of TUNEL-positive cells was also significantly increased, while the proportion of EdU-positive cells was decreased in MSTN-overexpressing CT-hSM-MPCs compared with control cells (Fig 3D and E & Appendix Fig S1). Moreover, increased ROS production was shown in MSTN-overexpressing CT-hSM-MPCs (Fig 3F). Decreased mRNA and protein levels of MYOG but significantly increased expression of NOX4 were found in MSTN-overexpressing CT-hSM-MPCs (Fig 3G and H). However, the protein levels of NOX2 were downregulated in MSTN-overexpressing CT-hSM-MPCs (Fig 3H).

**Figure 3.**
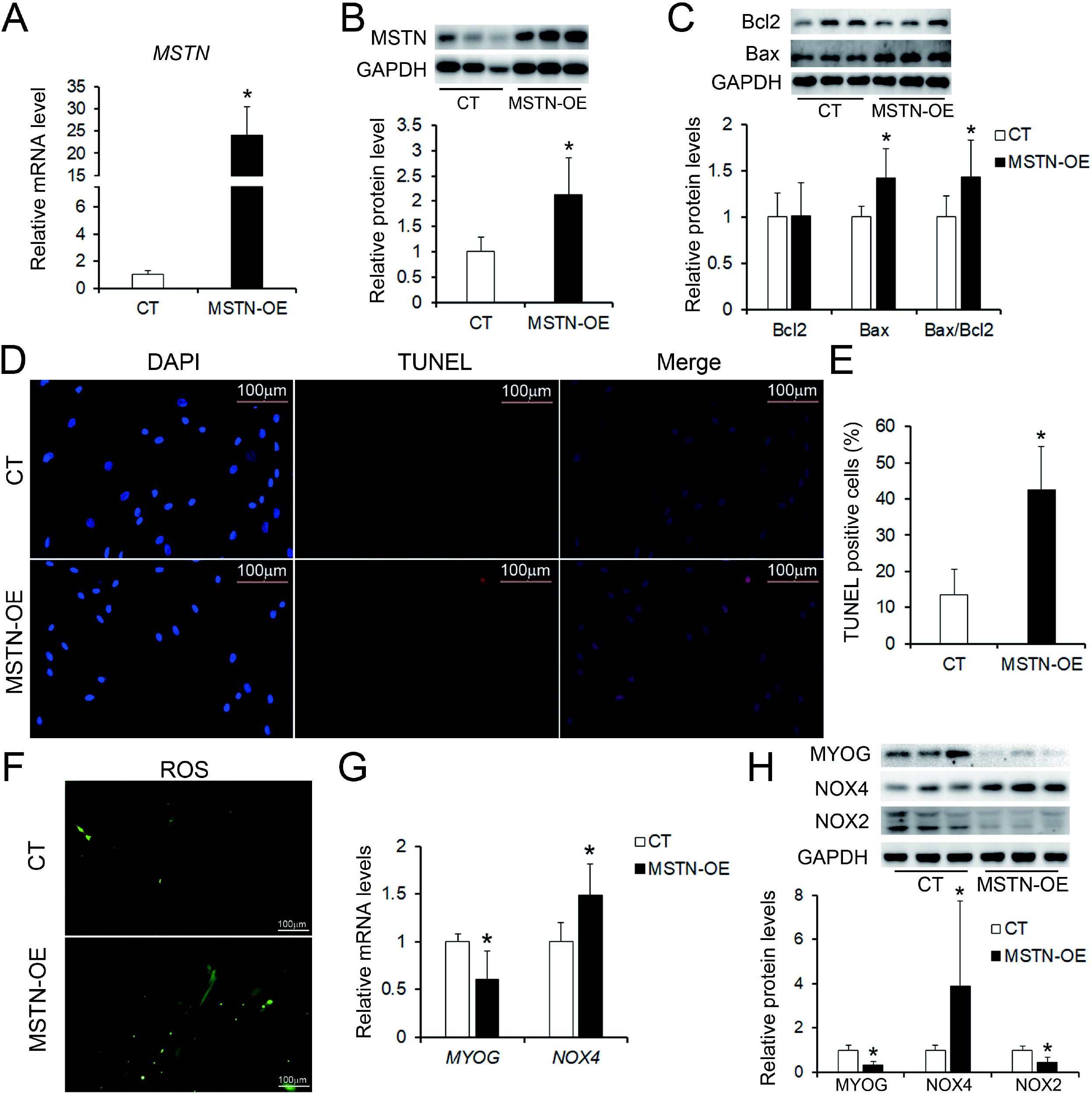
MSTN overexpression in CT-hSM-MPCs increase ROS release and apoptosis. A, B The mRNA and protein levels of MSTN in MSTN overexpressing CT-hSM-MPCs and control cells after transfection. C Western blotting analysis of apoptosis related proteins in MSTN overexpressing CT-hSM-MPCs and control cells. D, E TUNEL assay of MSTN overexpressing CT-hSM-MPCs and control cells and quantity evaluation of percentage of the TUNEL positive cells. F ROS production of MSTN overexpressing CT-hSM-MPCs and control cells. G, H RT-qPCR and Western blotting analysis of oxidative stress and myogenesis related proteins in MSTN overexpressing CT-hSM-MPCs and control cells. CT, empty pcDNA3.1(+) plasmids transfected CT-hSM-MPCs; MSTN-OE, pcDNA3.1 (+)-MSTN plasmids transfected CT-hSM-MPCs. n=9, *p<0.05.

### MSTN knockdown inhibits ROS production and apoptosis and induces myogenesis in IS-hSM-MPCs

We next silenced MSTN in IS-hSM-MPCs by siRNA. As shown in Fig 4A and B, NOX4, but not NOX2 or eNOS, was downregulated in MSTN-silenced IS-hSM-MPCs compared with NC siRNA-transfected IS-hSM-MPCs. Obvious decreases in ROS production were observed in two MSTN-silenced IS-hSM-MPC lines (Fig 4C). TUNEL assays also revealed decreased apoptosis after MSTN knockdown, as demonstrated by fewer TUNEL-positive cells in two MSTN-silenced IS-hSM-MPC lines than in their controls (Fig 4D and E). In addition, decreased levels of Bax protein and Bax/Bcl2 ratios were found in MSTN-silenced IS-hSM-MPCs compared with the control, while MYOG was upregulated and Bcl2 was unchanged in MSTN-silenced IS-hSM-MPCs (Fig 4F and G).

**Figure 4.**
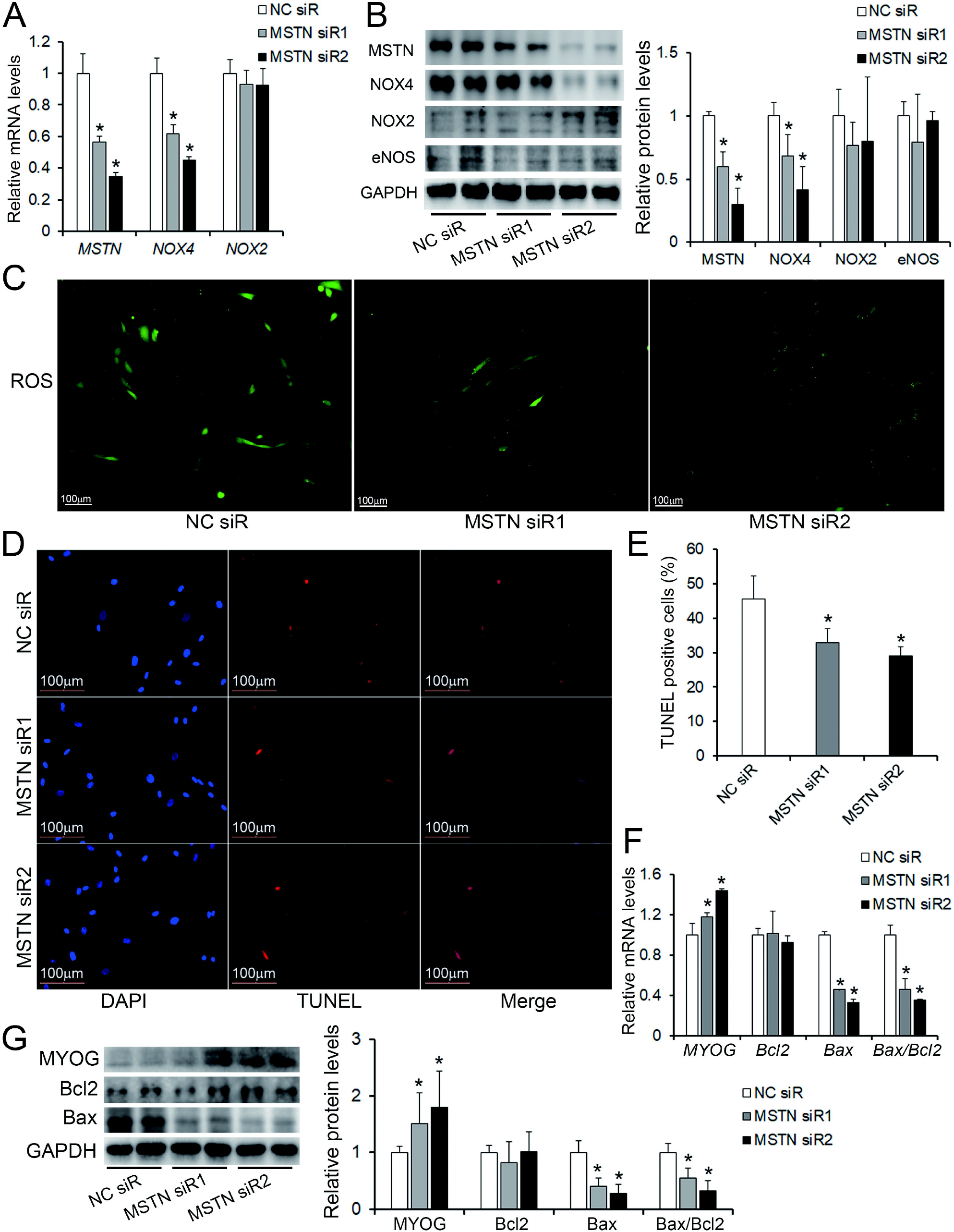
MSTN knockdown in IS-hSM-MPCs decrease ROS release and apoptosis. A, B The mRNA and protein levels of MSTN and oxidative stress related proteins in siRNA transfected IS-hSM-MPCs. C ROS production of siRNA transfected IS-hSM-MPCs. D, E TUNEL assay of siRNA transfected IS-hSM-MPCs and quantity evaluation of percentage of the TUNEL positive cells. F, G RT-qPCR and Western blotting analysis of apoptosis and myogenesis related proteins in siRNA transfected IS-hSM-MPCs. NC siR, NC siRNA transfected IS-hSM-MPCs; MSTN siR, MSTN siRNA transfected IS-hSM-MPCs. n=6, *p<0.05.

### MSTN overexpression in the PVM leads to scoliosis in mice

To test the roles of upregulated MSTN in the PVM in the development of IS, and to confirm the influence of differences in the two sides of the PVM on the occurrence of IS, we established a mouse model with right-side only or bilateral overexpression of MSTN specifically in the PVM by in situ injection of MSTN-AAV and control-AAV Three weeks after injection, thoracic vertebrae and nearby PVM tissues, as well as internal organs, of mice unilaterally injected with control virus were dissected for verification of PVM-specific overexpression (Fig EV3). Thirteen weeks after injection, a CT scan was carried out on the skeleton of the mice. We observed certain proportions of scoliosis occurrence, with a more than sixty percent occurrence found in bilateral PVM MSTN-overexpressing female (bi-MSTN-OE-F) and male (bi-MSTN-OE-M) mice, with no structural abnormality in the vertebral bodies (Fig 5A and B). Moreover, the incidence of scoliosis in the bi-MSTN-OE-F mice was higher than that in the bi-MSTN-OE-M mice, and the body weights of the bilateral PVM MSTN-overexpressing (bi-MSTN-OE) mice were decreased compared with that of the control (bi-CT) mice, which is similar to the disease characteristics observed clinically (Fig 5B and C). Similarly, a scoliosis phenotype was found in both unilateral PVM MSTN-overexpressing female (uni-MSTN-OE-F) and male (uni-MSTN-OE-M) mice, and the incidence rate of female mice was also higher than that of male mice (Fig 5D and E). It is worth noting that the incidence of scoliosis in bi-MSTN-OE mice was higher than that in unilateral PVM MSTN-overexpressing (uni-MSTN-OE) mice, which was less than fifty percent (Fig 5B and E). Moreover, minimal body weight difference between uni-MSTN-OE and uni-CT mice were observed (Fig EV4).

**Figure 5.**
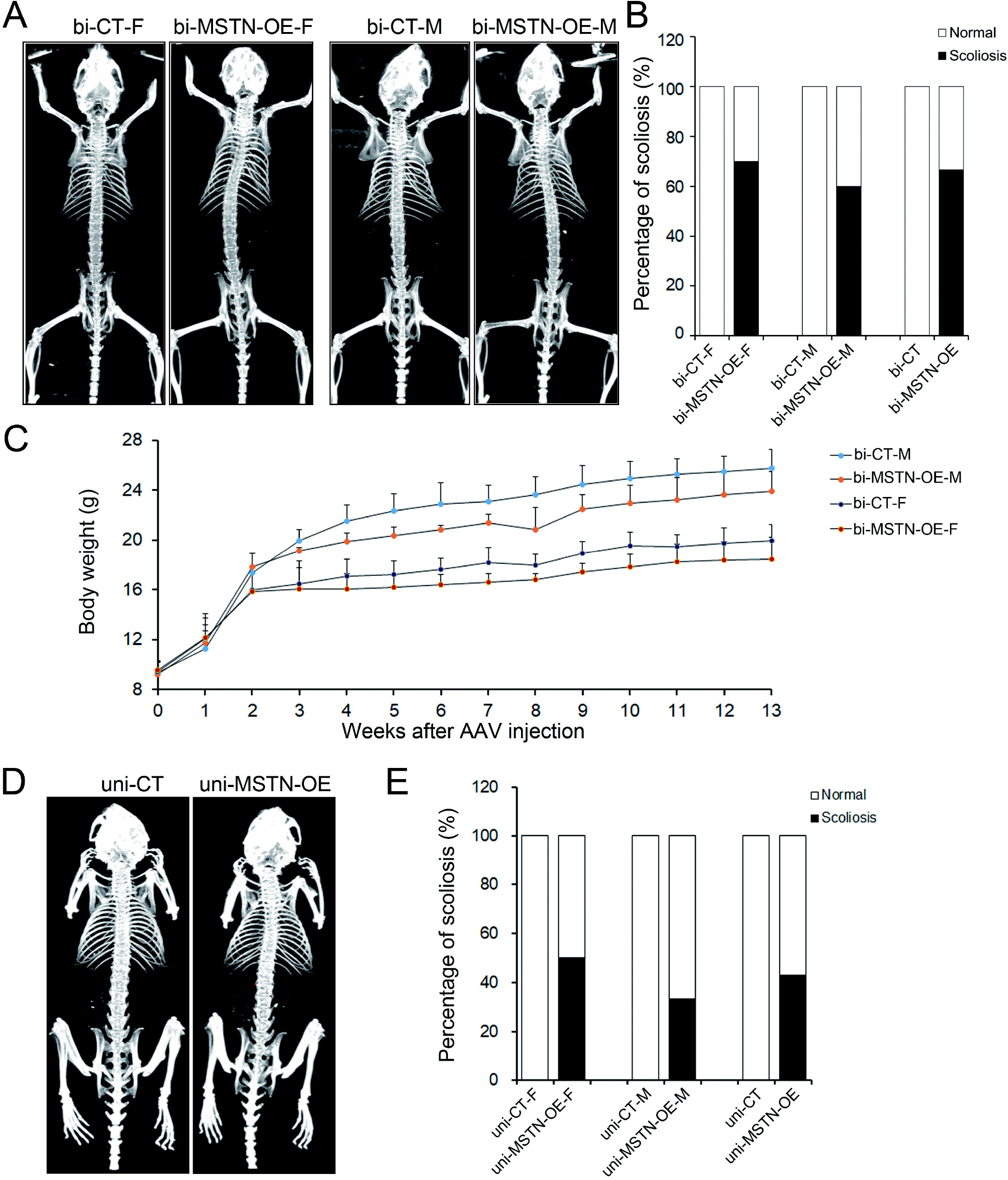
PVM overexpression of MSTN caused scoliosis phenotype in mice. A, B CT analysis of bilateral PVM AAV injected female and male mice and quantity evaluation of the percentage of scoliosis, n=15. C Growth chart of bilateral PVM AAV injected female and male mice after injection, n=15. D, E CT analysis of unilateral PVM AAV injected female and male mice and quantity evaluation of the percentage of scoliosis, n=6-7. bi-MSTN-OE-F, bilateral PVM MSTN overexpressing female mice; bi-MSTN-OE-M, bilateral PVM MSTN overexpressing male mice; bi-MSTN-OE, bilateral PVM MSTN overexpressing mice; bi-CT, bilateral PVM control mice; uni-MSTN-OE-F, unilateral PVM MSTN overexpressing female mice; uni-MSTN-OE-M, unilateral PVM MSTN overexpressing male mice; uni-MSTN-OE, unilateral PVM MSTN overexpressing mice; uni-CT, unilateral PVM control mice.

These results suggested that the occurrence of IS may be more related to abnormalities in PVM structure and function than to differences in the two sides of the PVM. Therefore, we next focused on the comparative analysis of bi-MSTN-OE and bi-CT mice.

### Elevated oxidative stress and mild muscle injury observed in bi-MSTN-OE mice

The expression levels of MSTN and the Flag tag protein in the PVM of the bi-MSTN-OE group were significantly higher than those of the bi-CT group (Fig 6A and B). The mRNA and protein levels of Bax and the Bax/Bcl2 ratio were significantly increased in the bi-MSTN-OE group, while the mRNA and protein levels of Myog were decreased in the bi-MSTN-OE group compared with the bi-CT group (Fig 6A and B). No significant changes in Bcl2 expression levels were observed between the two groups (Fig 6A-B). Although the hematoxylin-eosin staining results showed that the PVM of the bi-MSTN-OE group had only mild injury, the dihydroethidium (DHE) staining results showed significantly increased levels of ROS in the bi-MSTN-OE group (Fig 6C and F). The mRNA and protein levels of NOX4 in the PVM of the bi-MSTN-OE group were also increased compared with those of bi-CT mice, whereas the expression levels of NOX2 and eNOS between the two groups showed no significant differences (Fig 6D and E).

These results suggested that MSTN overexpression in the PVM can lead to the pathogenesis of IS, possibly elevating oxidative stress by upregulating NOX4 and further promoting apoptosis, as well as inhibiting myogenesis in the PVM.

## Discussion

The PVM is considered one of the most likely contributing factors for IS, and massive abnormalities such as necrosis, fibrosis, and fatty involution in PVM of IS patients have been reported (Jiang *et al.,* 2017; Peng *et al*, 2020). Although the exact underlying mechanism of severe muscle abnormities is unclear, scholars have revealed that mutated genetic factors related to PVM development, such as *LBX1* and *PAX3,* may be involved (Decourtye *et al*, 2022; Jennings *et al*, 2019; Qin *et al*, 2020; Wang *et al*, 2021; Zhu *et al*, 2015). In our previous study, compared with age-matched controls, the PVMs of IS patients showed elevated oxidative stress, especially increased ROS levels, which could induce severe muscle injury with abnormal myogenesis. In this study, increased apoptosis, impaired myogenesis and elevated oxidative stress were observed in IS-hSM-MPCs compared with CT-hSM-MPCs (Fig 1). It has been well demonstrated that hSM-MPCs support skeletal muscle development, growth, and regeneration (Wosczyna & Rando, 2018); thus, changes in IS-hSM-MPCs are responsible for PVM abnormities.

Oxidative stress is a biological phenomenon caused by an imbalance in redox systems and may involve either excessive production of reactive oxygen species or dysfunctional antioxidant enzymes (Saleem *et al*, 2020). Since increased expression of SOD1 (superoxide dismutase 1) and PGC1α (peroxisome proliferator activated receptor γ coactivator-1 α), which belong to the antioxidant system, was observed, we speculated that the production of reactive oxygen species was severely increased in the PVM of IS patients. Reactive oxygen species are produced not only by mitochondria but also by NOX (Specht *et al*, 2021; Xirouchaki *et al*, 2021). Our data showed that the expression level of NOX4, but not NOX2, which are the two main NOXs expressed in the PVM, was significantly increased in IS-hSM-MPCs (Fig 2). This suggested that NOX4 may play a role in the elevated oxidative stress in IS-hSM-MPCs. Even though mitochondrial abnormities were found in several studies, the function of mitochondria as it relates to increased ROS levels in IS patients is not yet known, and related investigations are still needed.

We further investigated the reasons for the abnormal changes in IS-hSM-MPCs related to elevated oxidative stress, and an upregulation of the negative muscle regulator MSTN was observed in both hSM-MPCs and the PVM of IS patients (Fig 2 and Fig EV2). MSTN, a murine TGF-beta family member also called growth/differentiation factor-8, includes a signal peptide, an N-terminal precursor peptide and a C-terminal mature peptide and acts by autocrine or paracrine signaling (McPherron *et al*, 1997). MSTN knockout animals showed a “double muscle character” with a significant increase in muscle mass, and natural mutation of MSTN in animals or humans also causes muscle hypertrophy (Gao *et al*, 2020). MSTN knockout can promote the proliferation and differentiation of hSM-

MPCs/myoblasts and allow them to resist apoptosis to improve their survival rate (Zhang *et al*, 2018a; Zhang *et al.,* 2018b). In addition, MSTN induced intracellular ROS release and upregulated NADPH oxidase in HK-2 tubular epithelial cells, while myostatin deletion prevented the increase in NOX-4 levels in the kidney (Butcher *et al*, 2018; Verzola *et al.,* 2020). Myostatin deficiency protects C2C12 cells (mouse myoblast cell line) from oxidative stress by inhibiting the intrinsic activation of apoptosis (Drysch *et al*, 2021). Meanwhile, our in vitro experiments verified that MSTN upregulation in CT-hSM-MPCs could impair myogenesis, induce apoptosis and promote the expression of NOX4 and release of ROS (Fig 3). Blocking MSTN upregulation in IS-hSM-MPCs through siRNA enhanced myogenesis, reduced apoptosis, and decreased NOX-4 and ROS levels (Fig 4). These results indicated that MSTN could cause elevated oxidative stress, together with the abnormal changes in hSM-MPCs and the PVM of IS patients, by regulating both NOX4 and myogenesis.

Next, we imitated PVM MSTN upregulation in mice and, surprisingly, observed a scoliosis phenotype in both bi-MSTN-OE and uni-MSTN-OE mice without vertebral abnormities (Fig 5). The incidence of scoliosis in MSTN-OE-F mice was higher than that in MSTN-OE-M mice, which was similar to the prevalence characteristics of IS being more common in girls than in boys (Cai *et al*, 2021; Chung *et al*, 2020). These results may be related to the innate muscle sexual dimorphism in both mice and humans that lower muscle mass in the arms and trunk were observed in females (Abe *et al*, 2021; O’Reilly *et al*, 2021). Moreover, the body weight of bi-MSTN-OE mice was decreased compared with that of bi-CT mice, which was also consistent with the phenomenon that IS patients have lower body mass (Cai *et al.,* 2021). Interestingly, the incidence of scoliosis observed in bi-MSTN-OE mice was much higher than that in uni-MSTN-OE mice. Therefore, we are more inclined to think that the development of IS due to abnormalities in PVM structure and function rather than concave-convex side differences. Unilateral PVM abnormalities have less effect on spinal stability than bilateral PVM abnormalities, which may be due to the compensation maintenance effect of normal PVMs on the other side.

In parallel with the in vitro results, MSTN upregulation resulted in severe oxidative stress, increased apoptosis, and abnormal myogenesis in mouse PVMs (Fig 6). These were the same PVM pathological features observed in IS patients (Li *et al.,* 2019a; Shahidi *et al*, 2021). Nevertheless, only mild necrosis was shown in bi-MSTN-OE mice, which may be associated with the slight scoliosis observed. Given that concave-convex side differences widen with the degree of curve, and that concavity is more affected in more severe IS patients, no significant changes between the concave and convex sides were found in these mice (Fig EV5). The skeletal growth of mice is fast during 2-5 weeks of age and slows during 5-8 weeks of age (Li *et al*, 2017). Therefore, we believe that earlier MSTN overexpression in the bilateral PVMs of mice would be more likely to affect spinal stability, since we only successfully in situ injected AAV into the PVMs of three-week-old mice, and three additional weeks following AAV injection were required for MSTN overexpression.

In conclusion, as shown in Fig 7, we found that the cause of oxidative stress elevation in IS patients could be MSTN-induced upregulation of NOX4, and MSTN plays important roles in the development of IS by regulating both oxidative stress and myogenesis. Our results may offer a great new and potential target for IS therapy, since MSTN deficiency leads to muscle hypertrophy with no severe adverse consequences (Gao *et al.,* 2020).

**Figure 6.**
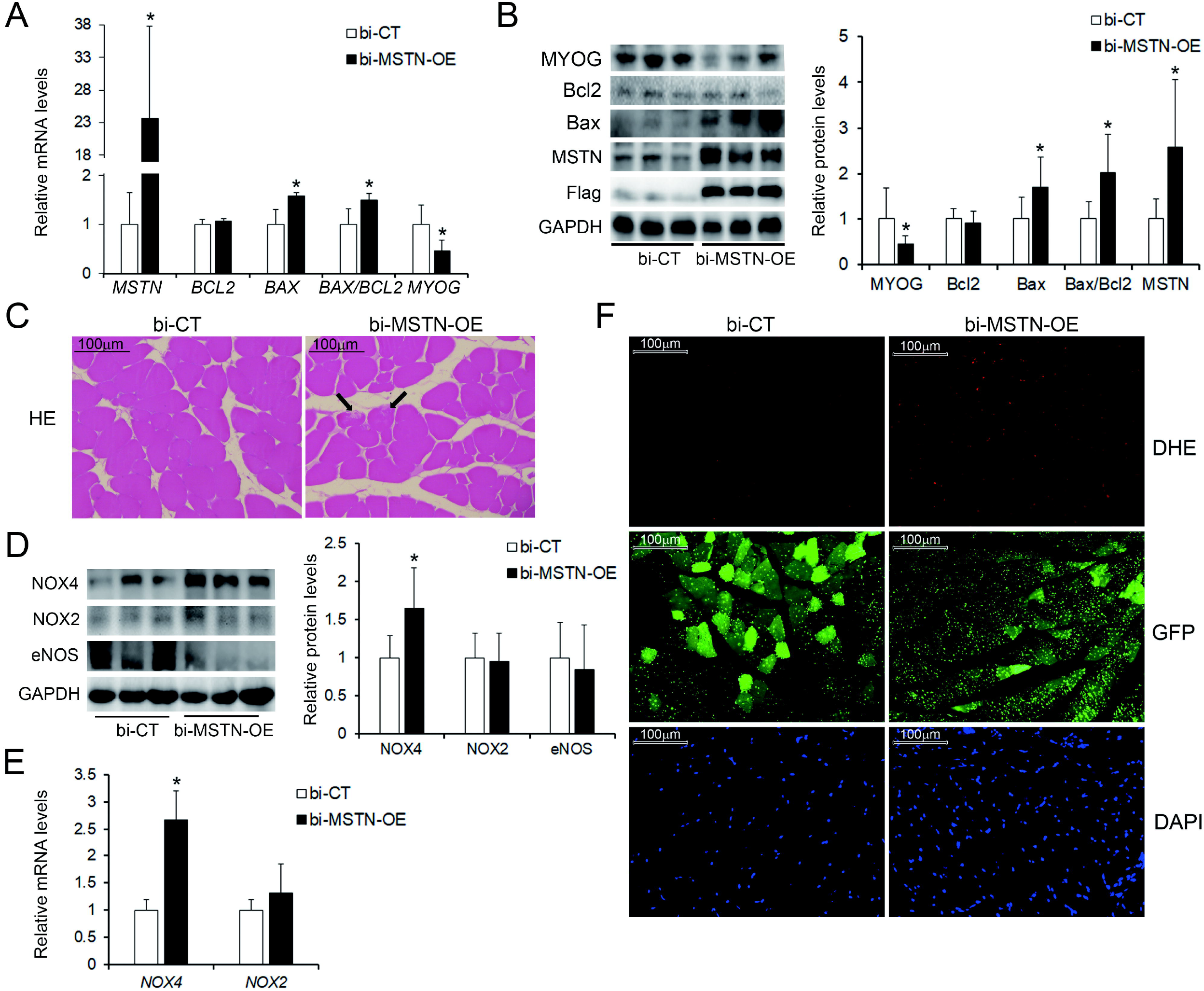
Alteration of PVM in bilateral PVM MSTN overexpressing mice. A, B The mRNA and protein levels of MSTN and apoptosis, myogenesis related proteins in PVM of bi-CT and bi-MSTN-OE mice. C hematoxylin-eosin stained PVM of bi-CT and bi-MSTN-OE mice, 200×, Black arrow indicates myofiber necrosis. D, E The mRNA and protein levels of oxidative stress related enzymes in PVM of bi-CT and bi-MSTN-OE mice. F DHE stained ROS production in PVM of bi-CT and bi-MSTN-OE mice. n=15. *p<0.05.

**Figure 7.**
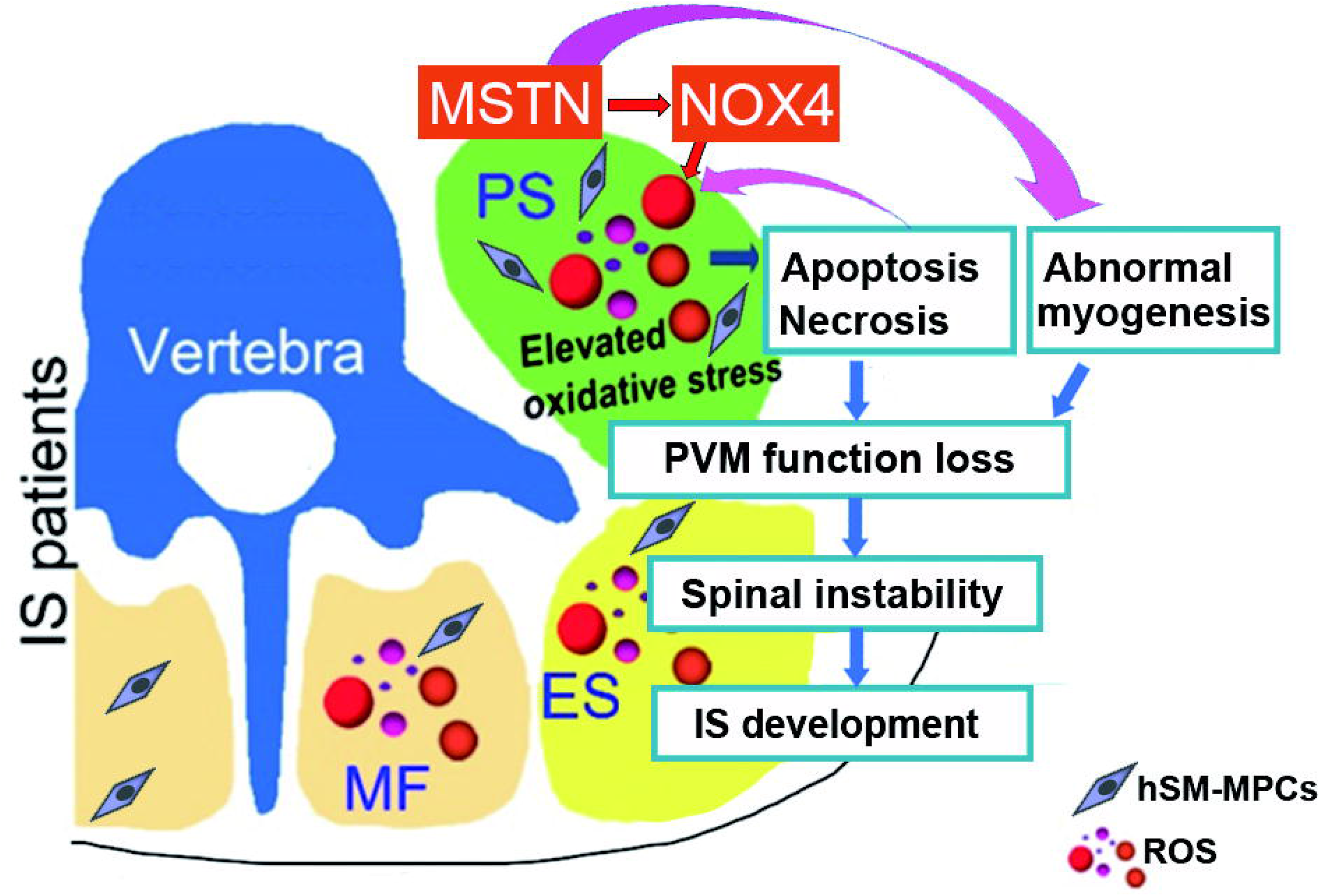
Possible roles of MSTN in the development of IS. Upregulated MSTN can increase NOX4 expression and elevate oxidative stress, promote apoptosis as well as inhibit myogenesis in PVM, which impact the function of PVM and further lead to the development of IS. PS, psoas; ES, erector spinae; MF, multifidus; ROS, reactive oxygen species; hSM-MPCs, primary skeletal muscle mesenchymal progenitor cells.

## Materials and Methods

### Patients, controls, and extraction of hSM-MPCs

The PVM tissues of 15 IS patients and 15 control subjects were collected carefully and safely during a posterior approach or by minimally invasive surgery in Xiangya Hospital (Table EV1); their hSM-MPCs were isolated enzymatically and cultured in growth medium at 37 °C/5% CO2 as described before (Li *et al.,* 2019a).

### TUNEL assay

Apoptotic cell death was detected using the One Step TUNEL Apoptosis Assay Kit with Cyanine 3 (Beyotime Biotechnology, Jiangsu, China) according to the manufacturer’s instructions. Briefly, cultured cells were fixed with 4% paraformaldehyde for 20 minutes (min), permeabilized with 0.3% Triton X-100 in PBS for 5 min at room temperature and subsequently incubated with TUNEL working solution for 1 hour at 37 °C. DAPI (Servicebio, Wuhan, China) was used to label the nuclei. TUNEL-positive cells were counted and reported as the percentage of TUNEL-positive cells.

### Immunofluorescent staining

Desmin immunofluorescent staining was used to evaluate the differentiation ability of extracted hSM-MPCs from controls and IS patients. Fixed cells were blocked for 30 min and then incubated with primary antibody (Santa Cruz Biotechnology, Dallas, USA) overnight at 4 °C. After extensive washing, sections were incubated with fluorescein-labeled anti-mouse immunoglobulin G (Abcam, Cambridge, UK) for 2 hours at room temperature. Cells were costained with DAPI.

### Measurement of superoxide anion and ROS assay

PVM tissues of mice were frozen in Tissue-Tek OCT embedding medium (Sakura Finetek, Tokyo, Japan) and cut into 8 μm thick sections. Then, 10 μM DHE (Beyotime Biotechnology) was used to evaluate superoxide levels in situ by incubating the slides in a dark chamber at 37 °C for 30 min (Tang *et al*, 2016). And hSM-MPCs were incubated with 10 μM DCFH-DA (ROS Assay Kit, Beyotime Biotechnology) at 37 °C for 20 min for ROS analyses according to the manufacturer’s instructions.

### Histological analysis and pathological evaluation

Fixed mouse PVM samples were embedded in paraffin and cut into 5 μm thick sections. Sections were stained with hematoxylin-eosin as previously described (Li *et al*, 2012). Pictures of 4-6 different fields per sample were taken under a Leica DM5000 B microscope equipped with a digital CCD (Leica Camera, Wetzlar, Germany).

### Western blotting

PVM tissues and hSM-MPCs were sonicated in RIPA buffer (Beyotime Biotechnology) to obtain whole-cell lysates. Western blotting analysis was performed as previously reported (Wang *et al*, 2016). Antibodies for eNOS were obtained from Cell Signaling Technologies (Danvers, MA), antibodies for NOX4 and NOX2 were purchased from Santa Cruz Biotechnology, antibodies for MYOG were from Servicebio, and the remaining antibodies were from Proteintech Group, Inc. (Chicago, USA). The expression levels of target proteins were quantified with Quantity One 1-D Analysis Software (Bio-Rad, Hercules, CA) and normalized to glyceraldehyde-3-phosphate dehydrogenase (GAPDH) in the same sample.

### Real-time PCR

Total RNA of hSM-MPCs and PVM tissues was extracted using TRIzol Reagent (CWBiotech, Beijing, China). Equal masses of RNA from each sample were reverse transcribed into cDNA using a HiFiScript cDNA Synthesis Kit (CWBiotech). Real-time PCR was performed with specific primers for the targeted genes by NovoStart^®^ Probe qPCR Super Mix (Novoprotein, Shanghai, China), and the transcriptional levels were quantified using GAPDH in the same sample as an internal control. The sequences of the primers used are described in Table EV2 or before (Li *et al.,* 2019a; Li *et al*, 2019b).

### RNA-seq and bioinformatic analysis

Total RNA was extracted from isolated hSM-MPCs using TRIzon Reagent (CWBiotech). RNA-seq library construction and high-throughput RNA sequencing was performed by Wuhan Genomic Institution (BGI HUADA, China) on a BGISEQ-500 high-throughput sequencer. Cluster analysis and Kyoto Encyclopedia of Genes and Genomes pathway analysis were performed according to a previously described method (Trapnell *et al*, 2012).

### Plasmids, siRNAs and transfection

The CDS of human *MSTN* from MSTN-pENTER plasmids (WZ Biosciences Inc., Shandong, China) was inserted into the pcDNA3.1+ vector by NheI/BamHI (New England Biolabs Inc., Ipswich, MA, USA) digestion. The positive plasmids were transfected into hSM-MPCs by Lipo8000 (Beyotime Biotechnology) following the user guide. NC siRNAs and siRNAs targeting human *MSTN* (the targeted sequences of MSTN siRNA1 and siRNA2 were TGACGATTATCACGCTACA and GTAGTAAAGGCCCAACTAT, respectively) were purchased from RiboBio Co. (Guangzhou, China). The siRNAs were transfected into hSM-MPCs by riboFECT CP transfection reagent (Ribobio Co.) following the user guide.

### Animal experiments and scoliosis phenotype analysis

Three-week-old wild-type C57BL/6 mice were obtained from Hunan SJA Laboratory Animal Co., Ltd. (Changsha, China) and housed in ventilated microisolator cages with free access to water for animal experiments. Body weight was measured weekly. For the bilateral PVM MSTN overexpression mouse model, we administered the HBAAV2/9-CMV-m-Mstn-3Xflag-ZsGreen and AAV2/9-CMV-3Xflag-ZsGreen control viruses, which were purchased from Hanbio Technology Ltd. (Shanghai, China), locally to both sides of the PVM around the thoracic vertebrae of three-week-old C57BL/6 mice. Fifteen mice were used for each group, with 4-5×10^10^ viral particles per injection and two to three injections on each side per mouse. For the unilateral PVM MSTN overexpression mouse model, we administered the two viruses locally to only the right PVM around the thoracic vertebrae of three-week-old C57BL/6 mice, with two to three injections per mouse, and seven mice were used for each group. Thirteen weeks after injection, mouse scoliosis phenotypes were examined using a NanoScan PET/CT (Mediso, Arlington, USA) at an energy of 50 kVp and an exposure of 186 μAs, with 480 projections per bed position after anesthetizing the mice with 4% chloral hydrate. Tomographic images were reconstructed to an isotropic voxel size of 250 μm using a filtered back-projection algorithm with a high-resolution Ram-Lak filter (Safaric Tepes *et al*, 2021). Interview Fusion software (Mediso) was used for CT image postprocessing. Thoracic vertebrae and nearby PVM tissues, as well as internal organs, were dissected after CT scans for further investigation. Animals were handled according to the Guidelines of the China Animal Welfare Legislation, as provided by the Committee on Ethics in the Care and Use of Laboratory Animals of Xiangya Hospital, Central South University.

## Statistical analysis

All results are expressed as the means ± standard deviation. Statistical significance was determined by analyzing the variance data using a Student’s *t test;* the gender difference was analyzed by the *x*^2^ test. Differences were considered statistically significant when p <0.05.

## Acknowledgements

The authors thank all the staf of the Department of Spine Surgery and Orthopaedics, Xiangya Hospital, Central South University for their dedicated assistance in patient sample collection. This work was supported by the National Natural Science Foundation of China (No.82072390) and the National Natural Science Foundation of Hunan Province, China (No.2020JJ4908).

## Author contributions

Jiong Li and Hongqi Zhang designed the study. Jiong Li performed most of the data collection, statistical analysis and data interpretation. Gang Xiang and Sihan He performed part of in vivo experiment. Guanteng Yang performed part of in vitro experiment. Chaofeng Guo and Mingxing Tang contributed to patient enrollment and follow-up. Jiong Li and Hongqi Zhang contributed to manuscript writing. All authors read and approved the final manuscript.

## Disclosure and competing interests statement

The authors declare that they have no competing interests.

## The Paper Explained

### PROBLEM

Idiopathic scoliosis (IS) is a disease with unknown etiology, characterized by spinal rotation asymmetry. However, paravertebral muscle (PVM) abnormalities play important roles in the pathogenesis of IS. Although our previous study found that elevated oxidative stress could inhibit myogenesis and promote apoptosis, resulting in PVM injury in IS patients, the underlying mechanism of oxidative stress generation is still unclear.

### RESULTS

Increased apoptosis, impaired myogenesis and elevated oxidative stress were found in primary skeletal muscle mesenchymal progenitor cells (hSM-MPCs) of IS patients, which are essential for the myogenesis process of vertebrate skeletal muscles. Through RNA-sequencing analysis between hSM-MPCs from controls and IS patients, 302 differentially expressed genes were observed. We further identified significantly upregulated myostatin (MSTN), which is a skeletal muscle negative regulator and can regulate reactive oxygen species release through NADPH oxidase 4, in IS hSM-MPCs. Overexpression of MSTN in hSM-MPCs from control patients increased the expression of NADPH oxidase 4, promoted reactive oxygen species production and apoptosis, and suppressed myogenesis. However, MSTN knockdown decreased the expression of NADPH oxidase 4, inhibited reactive oxygen species production and apoptosis, and enhanced myogenesis in hSM-MPCs from IS patients. In addition, overexpression of MSTN in the PVMs of mice induced markedly elevated oxidative stress and mild PVM injury, as well as scoliosis without abnormal vertebral structure.

### IMPACT

Altogether, our study suggested that abnormal PVM changes and accumulated oxidative stress in IS patients may result from upregulation of MSTN. Since overexpression of MSTN in the PVM can lead to scoliosis in vivo, it may contribute to the development of IS. Our results may offer a great new and potential target for IS therapy, since MSTN deficiency leads to muscle hypertrophy with no severe adverse consequences.

### For More Information

Information of MSTN can be found at OMIM Entry - * 601788 - MYOSTATIN; MSTN.

Information of other 24 controls and 33 IS patients (PVM tissues used for mRNA levels of MSTN) was mentioned in http://www.ijbs.com/v15p2584s1.pdf.

### List of abbreviations

IS, idiopathic scoliosis; AIS, adolescents with idiopathic scoliosis; PVM, paravertebral muscle; hSM-MPC, human skeletal muscle mesenchymal progenitor cell; IS-hSM-MPCs, hSM-MPCs from IS patients; CT-hSM-MPCs, hSM-MPCs from controls; Bax, BCL2-associated X protein; Bcl2, B cell leukemia/lymphoma 2; MYOG, myogenin; ROS, reactive oxygen species; eNOS, endothelial NO synthases; RNA-seq, RNA sequencing; MSTN, myostatin; NOX4, NADPH oxidase 4; NOX2, NADPH oxidase 2; bi-MSTN-OE-F, bilateral PVM MSTN overexpressing female mice; bi-MSTN-OE-M, bilateral PVM MSTN overexpressing male mice; bi-MSTN-OE, bilateral PVM MSTN overexpressing mice; bi-CT, bilateral PVM control mice; uni-MSTN-OE-F, unilateral PVM MSTN overexpressing female mice; uni-MSTN-OE-M, unilateral PVM MSTN overexpressing male mice; uni-MSTN-OE, unilateral PVM MSTN overexpressing mice; uni-CT, unilateral PVM control mice; DHE, dihydroethidium; min, minutes; GAPDH, glyceraldehyde-3-phosphate dehydrogenase.

### Ethics approval and consent to participate

This study has been conducted adhering to the principles of the Declaration of Helsinki II and approved by the medical ethics committee of Xiangya Hospital, Central South University (ethical code, 201703358). Written informed consent was obtained from all the study participants or their parent or guardian respectively.

### Data availability

The datasets used and/or analyzed during the current study are available from the corresponding author on reasonable request.

## Expanded View figure legends

Figure EV1 - KEGG analysis of 302 differentially expressed genes between hSM-MPCs from IS patients and controls. hSM-MPCs, human skeletal muscle mesenchymal progenitor cells.

Figure EV2 - Upregulated MSTN expression levels in PVM of IS patients compared with that of controls. Total RNA from PVM tissues of 39 controls and 48 IS patients were used for RT-qPCR, including PVM tissues from 24 controls and 33 IS patients mentioned in our previous study (Li *et al.,* 2019a). There were no differences of the sex and age between the 48 IS patients and 39 controls (Table EV1 and EV3). PVM, paravertebral muscles.

Figure EV3 - Verification of PVM MSTN overexpression at three weeks after control virus unilaterally injection in mice. PVM R, paravertebral muscles on the right side; PVM L, paravertebral muscles on the left side.

Figure EV4 - Growth chart of unilateral PVM MSTN overexpression mice after AAV injection.

Figure EV5 - Hematoxylin-eosin stained PVM tissue sections from bilateral PVM MSTN overexpression mice and the control mice.

**Table EV1.** Clinical data of study subjects for hSM-MPCs isolation.

| Group | Age | Gender | Lenke Type | Major curve<br>Cobb Angle (°) | Cause of the surgical intervention |
| --- | --- | --- | --- | --- | --- |
| C25 | 17 | M | - | - | Lumbar disc herniation |
| C26 | 21 | M | - | - | Lumbar disc herniation |
| C27 | 19 | F | - | - | Traumatic lumbar disc herniation |
| C28 | 11 | F | - | - | Lumbar disc herniation |
| C29 | 15 | M | - | - | Lumbar disc herniation |
| C30 | 16 | M | - | - | Lumbar disc herniation |
| C31 | 18 | F | - | - | Lumbar disc herniation |
| C32 | 25 | F | - | - | Lumbar disc herniation |
| C33 | 16 | M | - | - | Lumbar disc herniation |
| C34 | 15 | M | - | - | Traumatic lumbar disc herniation |
| C35 | 20 | F | - | - | Lumbar disc herniation |
| C36 | 18 | F | - | - | Traumatic lumbar disc herniation |
| C37 | 20 | F | - | - | Lumbar disc herniation |
| C38 | 21 | F | - | - | Lumbar disc herniation |
| C39 | 24 | M | - | - | Lumbar disc herniation |
| IS34 | 13 | M | 5 | 32.4 | Scoliosis |
| IS35 | 20 | F | 3 | 46.2 | Scoliosis |
| IS36 | 19 | F | 5 | 55.0 | Scoliosis |
| IS37 | 16 | F | 5 | 38.1 | Scoliosis |
| IS38 | 19 | M | 5 | 43.7 | Scoliosis |
| IS40 | 19 | F | 1 | 43.5 | Scoliosis |
| IS41 | 15 | M | 1 | 35.2 | Scoliosis |
| IS42 | 14 | F | 3 | 71.4 | Scoliosis |
| IS43 | 14 | F | 5 | 49.7 | Scoliosis |
| IS44 | 16 | F | 5 | 29.2 | Scoliosis |
| IS45 | 15 | M | 2 | 71.3 | Scoliosis |
| IS46 | 24 | F | 1 | 40.8 | Scoliosis |
| IS47 | 24 | F | 1 | 35.3 | Scoliosis |
| IS48 | 19 | F | 3 | 60.5 | Scoliosis |
| IS49 | 16 | M | 4 | 64.4 | Scoliosis |

C, control subjects; IS, idiopathic scoliosis.

**Table EV2.**
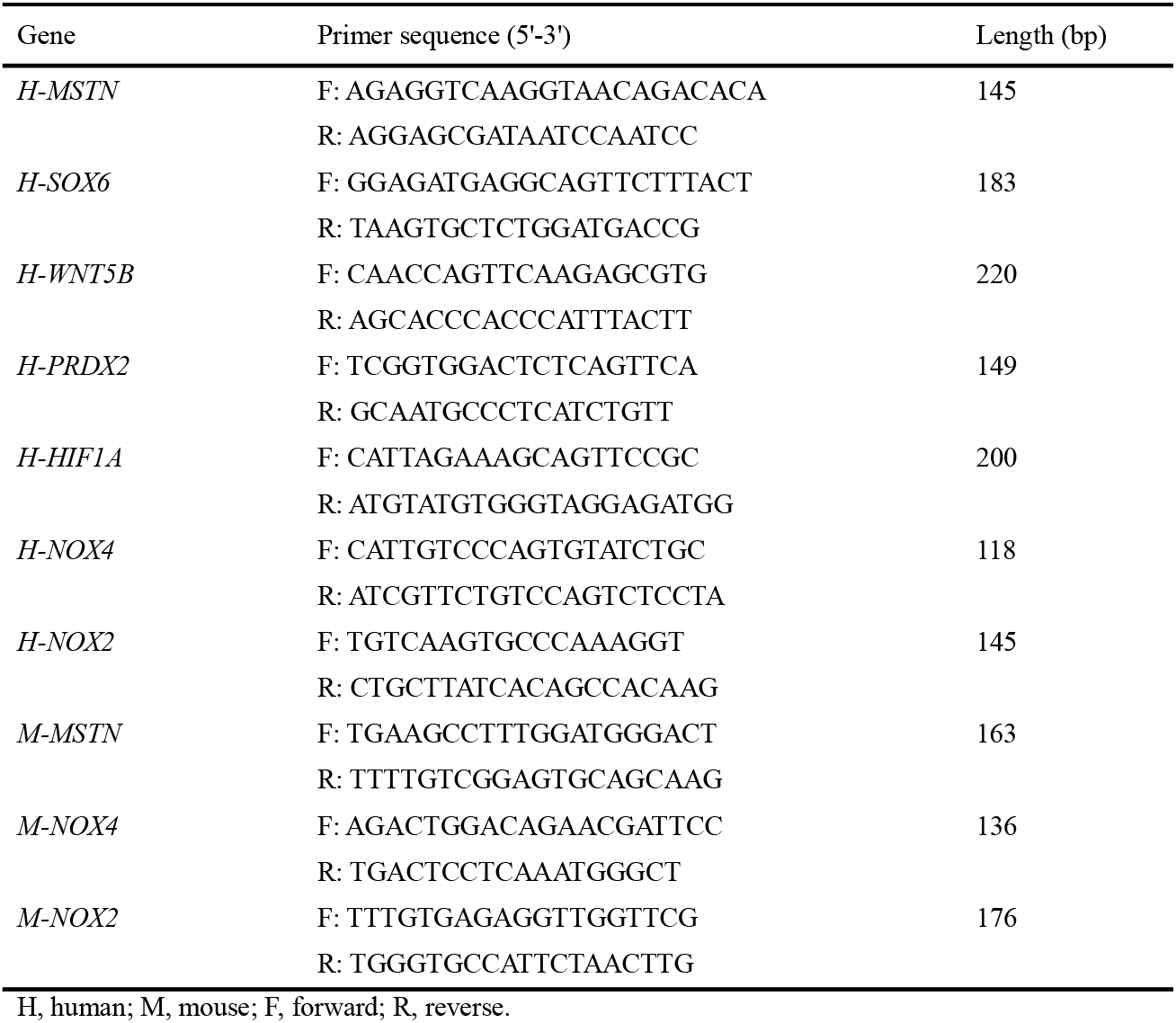
Primers sequences for genes.

**Table EV3.** Demographic of PVM study populations.

| Characteristics | CT *N*=39 | IS *N*=48 | *P* value |
| --- | --- | --- | --- |
| Age (years) | 17.68 ± 4.03 | 16.34 ± 3.25 | 0.091 |
| Gender | F 23 / M 16 | F 29 / M 19 | 1.000 |
| Major curve Cobb Angle (°) | - | 47.73 ± 14.79 | - |

CT, control group; IS, idiopathic scoliosis patients; F, female; M, male. Data was present as Mean ±
Standard Deviation.

## Notes

### Competing Interest Statement

The authors have declared no competing interest.

