## Supplementary figures and images for "Role of Paravertebral muscle myostatin upregulation in the development of idiopathic scoliosis"

### Fig EV1

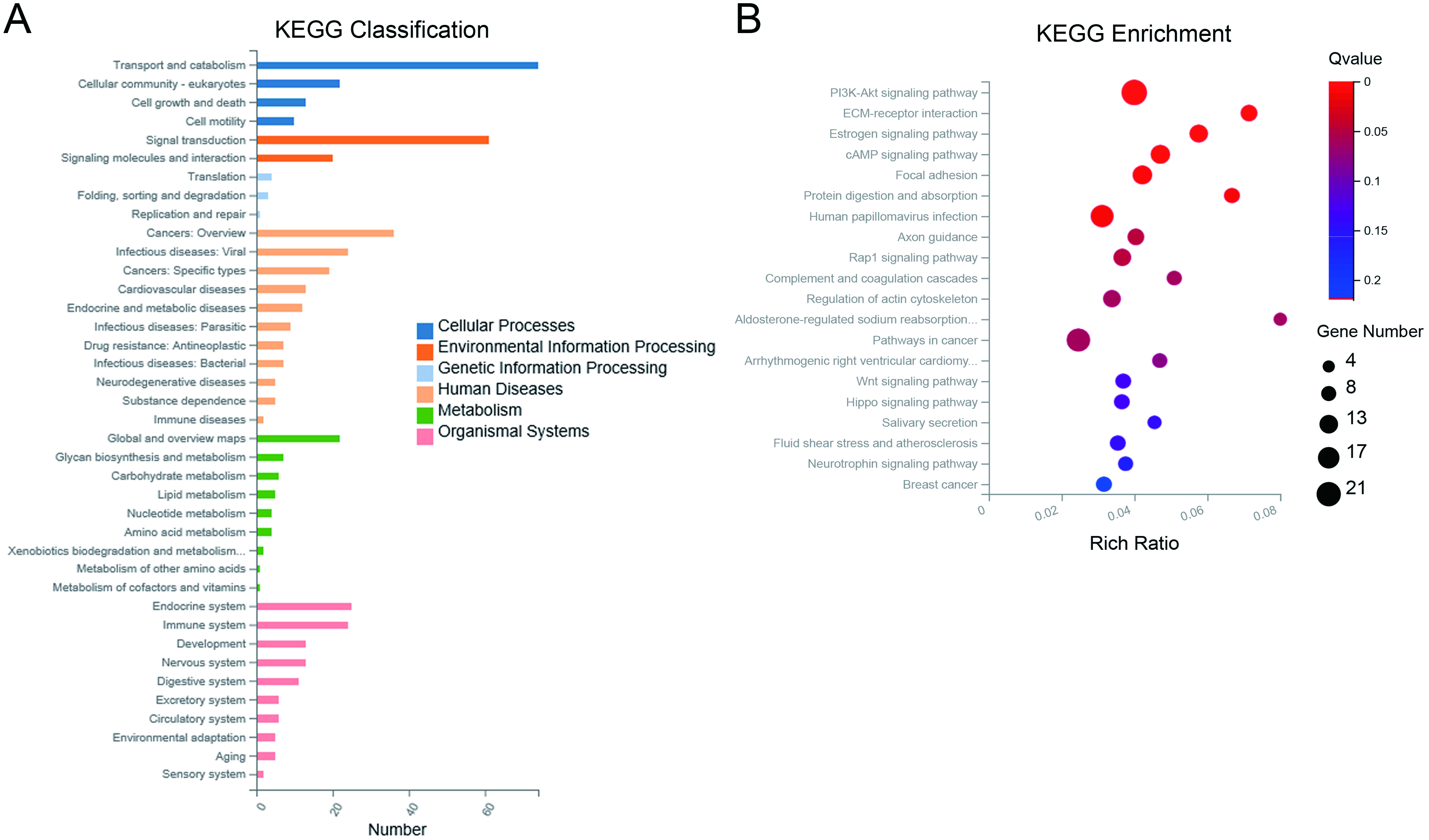

### Fig EV2

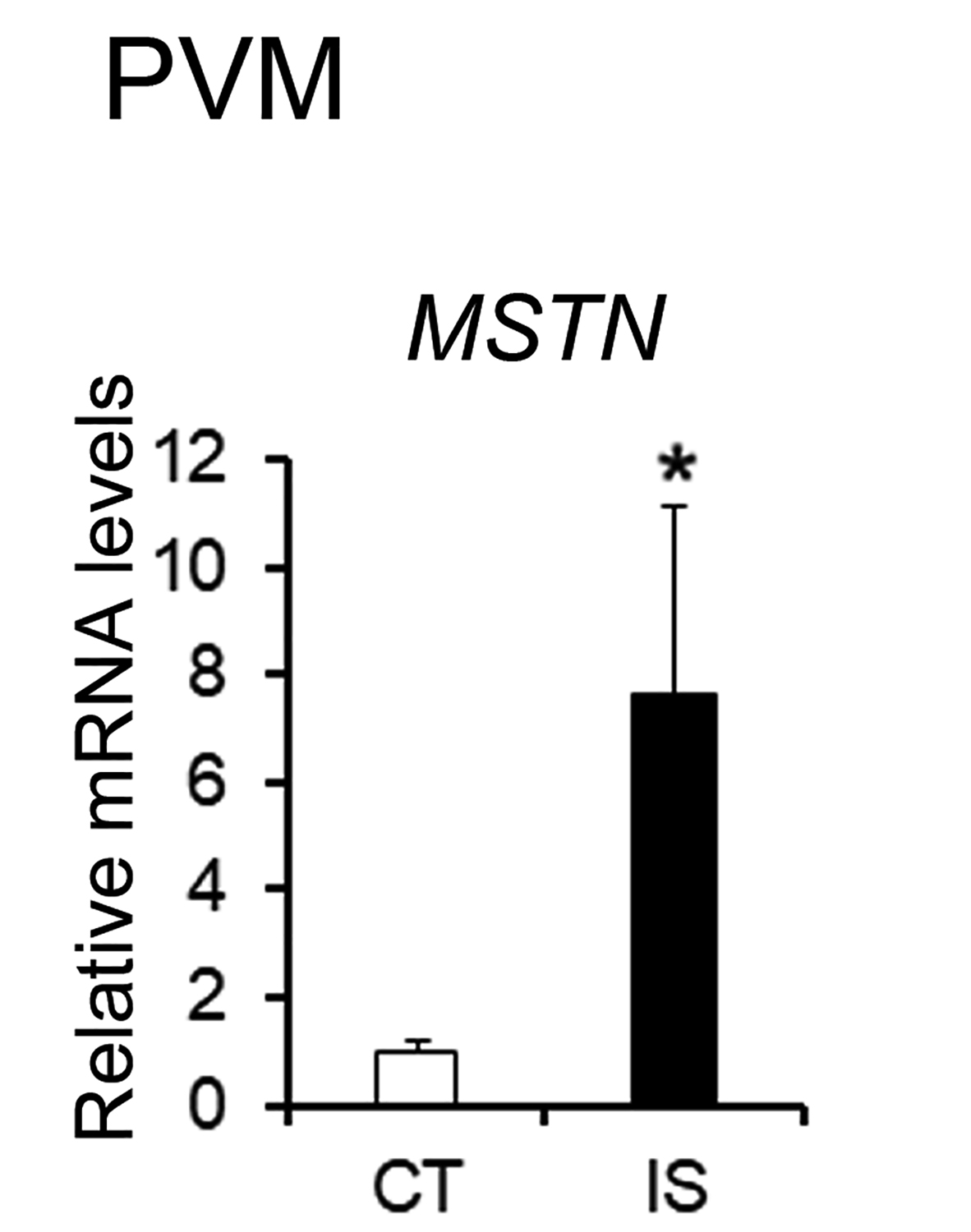

### Fig EV3

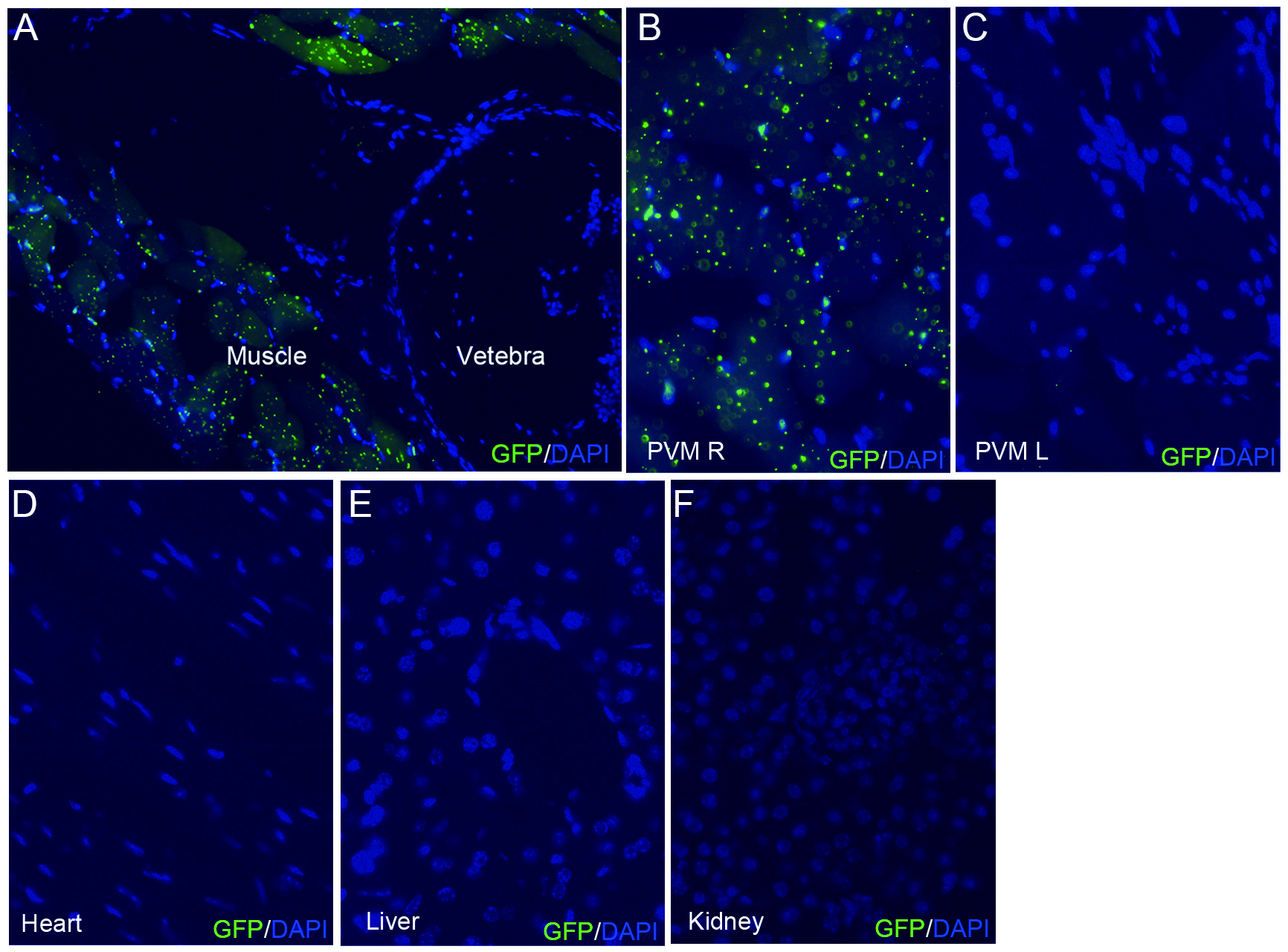

### Fig EV4

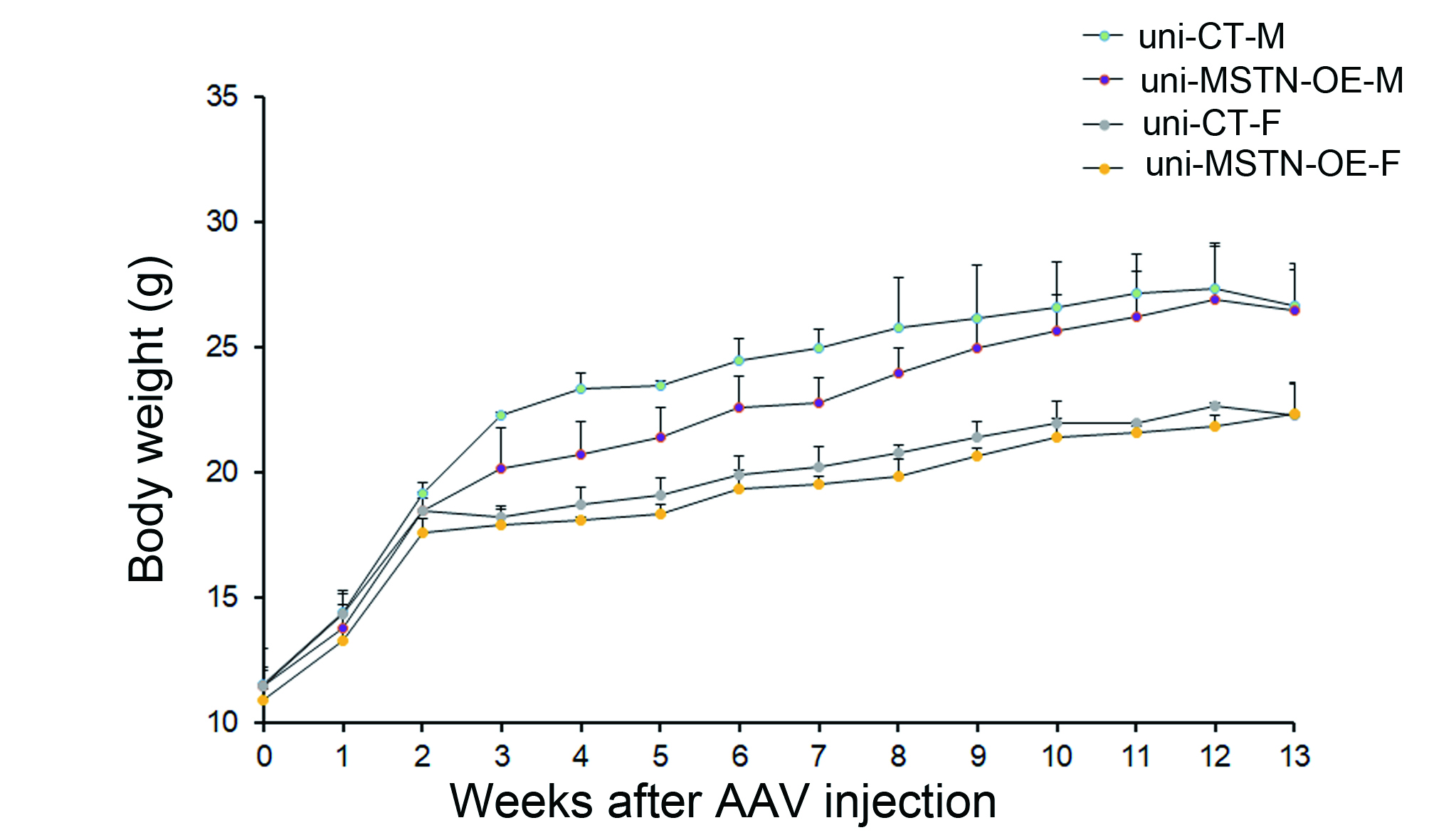

### Fig EV5

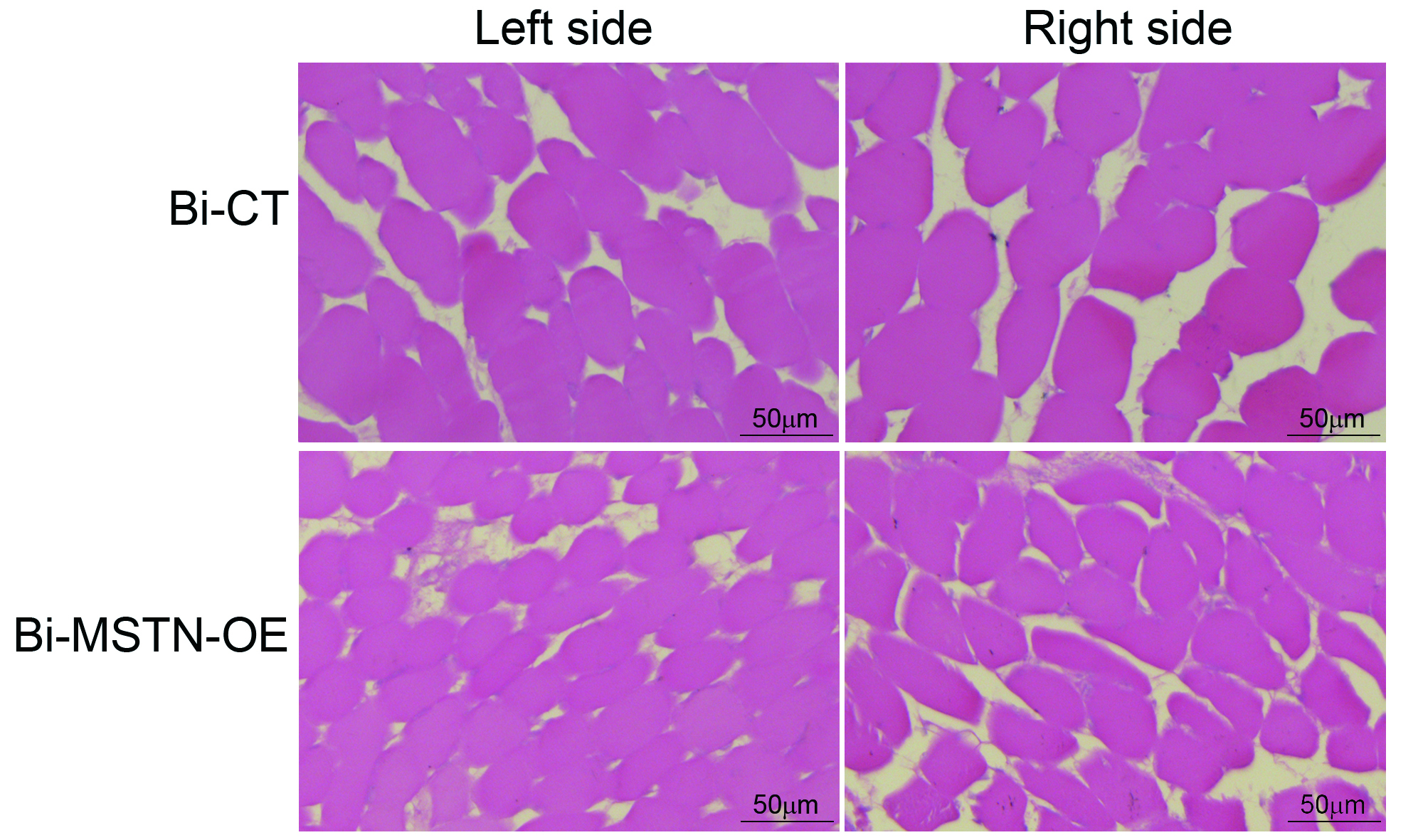
