## Appendix Fig S1 for "Role of Paravertebral muscle myostatin upregulation in the development of idiopathic scoliosis"

### Appendix PDF

| Contents | |
|---|---|
| Appendix Figure | Appendix Figure S1 |
| Appendix Supplementary Methods | EdU staining |

Appendix Figure

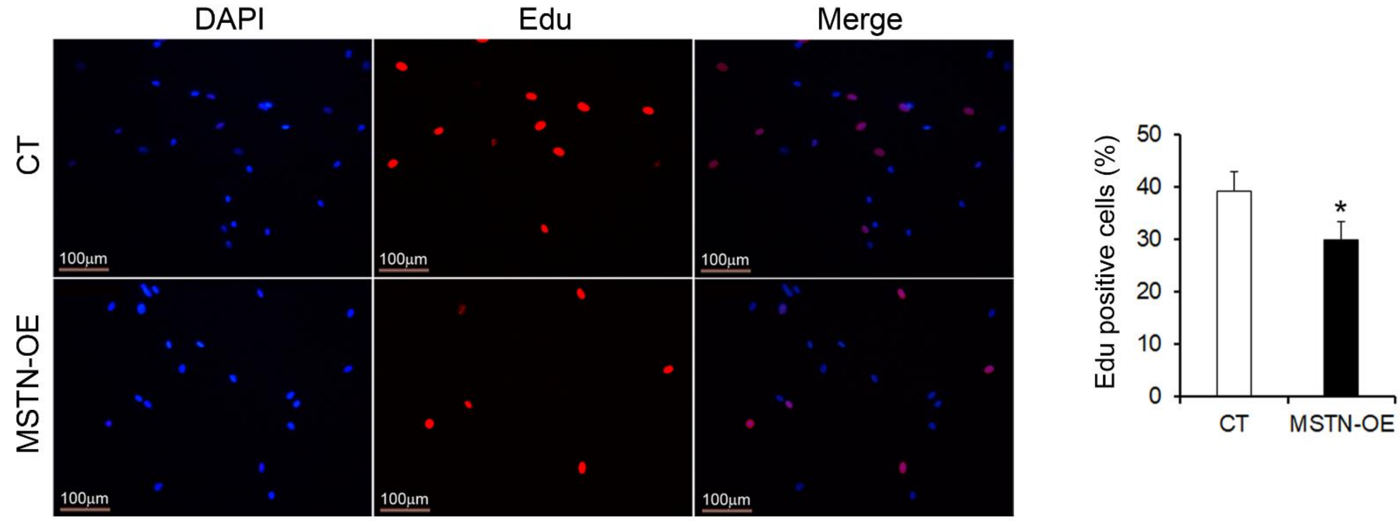


Appendix Figure S1 MSTN overexpression significantly decreased the proliferation of hSM-MPCs from controls. n=6. *p<0.05.

#### Appendix Supplementary Methods

##### EdU staining

EdU staining was conducted using the BeyoClick™ EdU Cell Proliferation Kit with Alexa Fluor 594 (Beyotime Biotechnology, Jiangsu, China, Cat. No. C00788S) according to the manufacturer's instructions. EdU was added into the medium until the final concentration was 10 μM, and the cells were incubated for 2 hours. After the incubation, the cells were washed with PBS to remove the DMEM and the free EdU probe and then fixed in 4% paraformaldehyde at RT for 15 min. After being co-stained with DAPI, the cells were observed under a Leica fluorescence microscope. EdU positive cells were counted at 5 different areas per sample and reported as a percentage of EdU-positive cells.
